# Organization of Area hV5/MT+ in Subjects with Homonymous Visual Field Defects

**DOI:** 10.1101/241836

**Authors:** Amalia Papanikolaou, Georgios A. Keliris, T. Dorina Papageorgiou, Ulrich Schiefer, Nikos K. Logothetis, Stelios M. Smirnakis

## Abstract

Damage to the primary visual cortex (V1) leads to a visual field loss (scotoma) in the retinotopically corresponding part of the visual field. Nonetheless, a small amount of residual visual sensitivity persists within the blind field. This residual capacity has been linked to activity observed in the middle temporal area complex (V5/MT+). However, it remains unknown whether the organization of hV5/MT+ changes following V1 lesions. We studied the organization of area hV5/MT+ of five patients with dense homonymous defects in a quadrant of the visual field as a result of partial V1+ or optic radiation lesions. To do so, we developed a new method, which models the boundaries of population receptive fields directly from the BOLD signal of each voxel in the visual cortex. We found responses in hV5/MT+ arising inside the scotoma for all patients and identified two possible sources of activation: 1) responses might originate from partially lesioned parts of area V1 corresponding to the scotoma, and 2) responses can also originate independent of area V1 input suggesting the existence of functional V1-bypassing pathways. Apparently, visually driven activity observed in hV5/MT+ is not sufficient to mediate conscious vision. More surprisingly, visually driven activity in corresponding regions of V1 and early extrastriate areas including hV5/MT+ did not guarantee visual perception in the group of patients with post-geniculate lesions that we examined. This suggests that the fine coordination of visual activity patterns across visual areas may be an important determinant of whether visual perception persists following visual cortical lesions.

## Introduction

The primary visual cortex (area V1) is considered to be the chief relayer of visual information to higher (extrastriate) visual areas. Lesions of V1 or its inputs lead to loss of conscious vision in specific parts of the contralateral visual hemifield (scotoma). The extent of the visual field defect corresponds retinotopically to the cortical region affected. Several studies have examined whether the adult V1 is able to reorganize following injury of the visual pathways (Wandell and Smirnakis, 2009). Less is known about the organization of extrastriate visual areas following chronic deprivation of V1 input (Goebel et al., 2001; Schmid et al., 2009).

Area V5/MT+ is of particular interest as it receives direct V1 input, it is retinotopically organized and it has been associated with the phenomenon of subconscious visual perception, called “blindsight” (Poppel et al., 1973; Weiskrantz et al., 1974). Experiments in macaque and New World marmoset monkeys showed that a significant proportion of V5/MT cells remain visually responsive in the absence of area V1 input (Bruce et al., 1986; Rodman et al., 1989; Maunsell et al., 1990; Rodman et al., 1990; Girard et al., 1992; Rosa et al., 2000; Schmid et al., 2010). In addition, Rosa et al. (Rosa et al., 2000) showed that many area MT neurons have ectopic receptive fields responding to the visual field surrounding the scotoma, suggesting reorganization. In contrast, experiments on New World owl monkeys showed that V5/MT depends entirely on V1 for visual activation (Kaas and Krubitzer, 1992; Collins et al., 2003; Collins et al., 2005). The basis of this discrepancy is not yet understood, and may be due to a species related difference.

In humans, the middle temporal complex (hV5/MT+) has been shown to be modulated by motion stimuli presented inside the scotoma following V1 lesions. Visual-motion related activity was observed in hV5/MT+ when moving stimuli were presented inside the blind visual field of a well-studied patient (G.Y.) with extensive area V1 injury (Barbur et al., 1993; ffytche et al., 1996; Zeki and Ffytche, 1998; Morland et al., 2004), a patient with homonymous hemianopia and Riddoch syndrome (Schoenfeld et al., 2002), a patient with bilateral damage to the gray matter of V1 (Bridge et al., 2010) and seven patients with chronic unilateral damage to V1 (Ajina et al., 2015). However, it is not known how the organization of area hV5/MT+ changes following V1 injury. Studying the patterns of activation in extrastriate visual areas following V1 lesions is important to understand the pathways that mediate residual visual behavior (“blindsight”) and to in the effort to guide future visual rehabilitation.

Here we studied the organization of hV5/MT+ area in five patients with homonymous visual field defects due to partial lesions of area V1 or the optic radiation. To do so, we developed a new method, which models the boundaries of population receptive fields (pRF) directly from the BOLD signal of each voxel in the visual cortex (Methods). The development of a new methodology was necessary here in order to minimize inaccuracies of pRF estimation in hV5/MT+ that are observed in healthy subjects with simulated visual field scotomas (Papanikolaou et al., 2015) when using existing pRF mapping methods (Dumoulin and Wandell, 2008; Lee et al., 2013).

Using this approach, we found that hV5/MT+ of the ipsilesional hemisphere does respond to visual stimuli presented within the scotoma. In 4/5 patients, it is possible that these responses originate from a partially damaged part of area V1 that gets activated by stimulus presented within the visual field scotoma. Nevertheless, this activity is apparently not sufficient to mediate conscious vision. In 2/5 patients (one patient showed both patterns), hV5/MT+ responses arise despite lack of significant corresponding V1 activation suggesting that it depends on the existence of functional V1-bypassing pathways. Cortico-cortical connective field modeling between areas V1 and hV5/MT+ supports these findings. Moreover, we found increased pRF sizes in hV5/MT+ of both hemispheres in three out of five patients suggesting the in some cases hV5/MT+ may undergo significant reorganization.

## Materials and Methods

### Subjects

*Patients*. Five subjects (27–64 years old, 3 females) with visual cortical lesions participated in our study. Four patients were recruited at the Center for Ophthalmology of the University Clinic in Tuebingen and one at the Core for Advanced MR Imaging of the Baylor College of Medicine (BCM). Four of the participants suffered from homonymous visual field deficits as a result of ischemic or hemorrhagic stroke 7–10 years before they enrolled in our study. One patient sustained an ischemic stroke 0.5 years prior to study recruitment (Table 1). The extent and location of the injury was confirmed by MRI anatomical acquisition (see below).

### Controls

Five healthy subjects (22–65 years old, 4 females) were recruited as controls. All subjects, patients and controls, had normal or corrected-to-normal visual acuity. The experiments were approved by the Ethical Committee of the Medical Faculty of the University of Tuebingen, and the IRB committee of BCM.

### Perimetric visual field tests

All patients underwent a Humphrey type (10–2) visual field test (Beck et al., 1985; Trope and Britton, 1987), with a (low photopic) background luminance level of 10 cd/m^2^. We present the Humphrey pattern deviation plots for all patients in Figure 2B. Patients P1, P2, P4 and P5 underwent additionally a binocular semi-automated 90° kinetic perimetry obtained with the OCTOPUS 101-perimeter (HAAG-STREIT, Koeniz, Switzerland) (Hardiess et al., 2010) which verified the visual field defects (Figure 2C).

### Data acquisition and preprocessing

Functional and structural MRI experiments were performed at the Max Planck Institute for Biological Cybernetics, Tuebingen, Germany and at the Core for Advanced MR Imaging of BCM, using a 3.0 Tesla high-speed echo-planar imaging device (Trio, Siemens Ltd., Erlangen, Germany) with a quadrature head coil. At least two T1-weighted anatomical volumes were acquired for each subject with a three-dimensional magnetization prepared rapid acquisition gradient echo (T1 MPRAGE scan) and averaged following alignment to increase signal to noise ratio (matrix size=256×256, voxel size=1×1×1 mm^3^, 176 partitions, flip angle=9°, TR=1900 ms, TE=2.26 ms, TI=900 ms). At Tuebingen, blood oxygen level dependent (BOLD) image volumes were acquired using gradient echo sequences of 28 contiguous 3 mm-thick slices covering the entire brain (repetition time TR=2,000 ms, echo time TE=40 ms, matrix size=64×64, voxel size=3×3×3 mm^3^, flip angle=90°). Functional image (echo planar) acquisition at BCM consisted of 29, 3.6mm-thick slices covering the entire brain (TR=2,000 ms, TE=30 ms, matrix size=64×64, voxel size=3.46×3.46×3.6 mm^3^, flip angle=90°; 200 volumes).

At least 5 functional scans were acquired for each subject, consisting of 195 image volumes, the first 3 of which were discarded. The functional images were corrected for motion in between and within scans (Nestares and Heeger, 2000). Subsequently, fMRI data were averaged across scans. The functional images were aligned to the high-resolution anatomical volume using a mutual information method (Maes et al., 1997) where the resampled time series values in the volume are spatially interpolated relative to the nearest functional voxels. Preprocessing steps were performed in MATLAB using the VISTASOFT toolbox (https://github.com/vistalab/vistasoft).

### Stimuli

Subjects were presented with moving square-checkerboard bars (100% contrast) through MRI compatible digital goggles (VisuaStim, Resonance Technology Company, Inc, Northridge, CA, USA; 30° horizontal and 22.5° vertical field of view, 800x600 resolution, min luminance=0.3cd/m^2^, max luminance=12.2cd/m^2^). The stimulus was presented within a circular aperture with a radius of 11.25° around the fixation point. The bar width was 1.875° and travelled sequentially in 8 different directions, moving by a step half of its size (0.9375°) every image volume acquisition (TR=2 seconds). Stimuli were generated using Psychtoolbox (http://psychtoolbox.org/) (Brainard, 1997) and an open toolbox (VISTADISP, https://github.com/vistalab/vistadisp) in MATLAB (The Mathworks, Inc.). The subjects’ task was to fixate on a small dot in the center of the screen (radius: 0.0375°; 2 pixels) and respond to the color change (red to green) by pressing a button. The color was changing randomly with a frequency of one every 6.25 seconds. An infrared eye tracker was used to record eye movements (iView XTM, SensoMotoric Instruments GmbH). The eye movement traces of patients P1, P2, P4 and P5 are shown in Figure S5. One patient (P3) was not eyetracked. However, his accuracy at the challenging fixation task was always more than 80% suggesting that the subject could maintain fixation.

Control subjects were asked to participate for a second session during which an isoluminant mask was placed in the left superior quadrant of the visual field, simulating a left upper quadrantanopia (“artificial scotoma” or AS). All other stimulus' parameters stayed the same.

### Population receptive field mapping

*<u>Direct-fit population receptive field (pRF) method:</u>* To define the borders between visual areas we derived pRF estimates using a direct-fit pRF method. This method has been described in detail before in (Dumoulin and Wandell, 2008). In short, the implementation of the pRF model is a circularly symmetric Gaussian receptive field in visual space. The center and radius of the pRF are estimated by fitting the BOLD signal responses to estimated responses elicited by convolving the model with the moving bar stimuli.

*<u>Topography-based pRF method:</u>* We also derived pRF estimates using a topography-based pRF method. We have described this method in detail in (Lee et al., 2013; Lee et al., 2015). In contrast to direct-fit methods (Dumoulin and Wandell, 2008), the topography-based pRF method does not assume a priori the pRF shape (e.g. circular) and thus is useful for studies of reorganization where the actual pRF shape cannot be anticipated. We used this method to confirm the location of areas V1 and hV5/MT+. We retained only those voxels in these visual areas, for which the topography explained more than 12% of the variance. This threshold was set after measuring the mean explained variance in a non-visually responsive area (6% ± 2%) and setting the value of the threshold at 3 standard deviations above the mean. Subsequently we used a new method to map the pRF boundaries for all voxels in the selected region of interest (ROI).

*<u>pRF boundary mapping method:</u>* We estimated the boundaries of the pRF directly from the BOLD time series of each voxel in the visual cortex. The pRF boundaries were identified by marking the location in the visual field where the BOLD activity starts to rise above a visual response threshold separately for each bar direction. The following steps were taken in order to estimate this: 1) A deconvolution method was applied to the BOLD time series of each voxel in order to estimate the underlying neural response of the voxel as the stimulus is presented at each visual field location. To do so, the BOLD time series of each voxel were averaged across scans (5–8 scans) to increase the signal to noise ratio. The averaged signal was further smoothed using locally weighted linear regression (lowess method in MATLAB) in order to avoid outliers that can be amplified after deconvolution. We then, applied Fourier deconvolution to remove the hemodynamic response function from the data. 2) A baseline was calculated from the deconvolved signal of each voxel: First, we calculated the troughs of the deconvolved BOLD time series using function *findpeaks* (MATLAB). A minimum trough distance was set according to the stimulus duration for each bar direction. This way, only the troughs that correspond to each bar direction are identified and averaged to estimate a general baseline. Then we estimated the noise level as the standard deviation of the signal when the bar stimulus is located in non-visually responsive locations of the visual field (e.g. >7° in the ipsilateral visual field). The visual response threshold is then estimated as the baseline plus 3 standard deviations of noise level. Using a lower threshold (baseline plus 1–2 standard deviations of noise level) does not significantly change our results. If the maximum response of a voxel was lower than the visual response threshold it was excluded from the analysis. 3) The deconvolved BOLD signal of each voxel was then considered separately for each bar direction and a Gaussian model was fit to the data. We chose the best model between a one-term and a two-term Gaussian mixture model using the Akaike’s information criterion (Burnham et al., 2002). The pRF boundaries were estimated by marking the location in the visual space at the time the fitted signal rises above threshold for each bar direction. This forms an octagon (since there are 8 different bar directions) in visual space, which represents the pRF. We only retained well-defined pRFs for which the Gaussian model explained more than 60% of the variance for each bar direction. A schematic representation of the method is shown in Figure S1. By taking into account only the activity rise when the stimulus approaches the border of the voxel receptive field from the outside, this method avoids errors in the pRF estimation that result from persistent hemodynamic activity that occurs when the bar stimulus moves from seeing to non-seeing locations of the visual field (Papanikolaou et al., 2015).

The pRF center is estimated as the center of “mass” of the octagon and the pRF size as the area of the octagon. The pRF amplitude of each voxel is estimated as the mean peak amplitude of the fitted Gaussian for each bar direction. The average pRF amplitude of an area in the lesioned hemisphere is normalized by the average pRF amplitude from the same area on the contra-lesional hemisphere.

#### Visual field coverage density maps

The visual field coverage density maps define the locations within the visual field that are covered by the pRFs of voxels within a region of interest (ROI) in the cortex. To estimate this we plot at each visual field location the number of the pRFs that cover this location (color map). The pRF centers (estimated as described above) across all voxels within the ROI are overlaid as grey dots.

#### Connective field (CF) modeling

CF parameters are estimated from the BOLD time-series using the CF modeling method described by Haak et al. (2013). Specifically, the fMRI response of each voxel in hV5/MT+ is predicted using a 2-dimensional circular Gaussian connective field model, folded to follow the cortical surface of V1. The CF of a voxel is defined by two parameters, the connective field position and the Gaussian spread across the V1 surface. A time-series prediction is then calculated by weighting the CF with the BOLD time-series. The optimal CF parameters are found by minimizing the residual sum of squares between the model’s time-series prediction and the observed time-series. Best models were retained if the explained variance in the observed fMRI time-series exceeded 12.5%. This threshold was set after measuring the mean explained variance in a non-visually responsive area (3.2% ± 3.1%) and setting the value of the threshold at 3 standard deviations above the mean. Our results are not sensitive to the specific choice of threshold over a range of 8–15%.

Because CF preferred locations in V1 cortical surface are associated with preferred visual field positions during the area V1 pRF mapping, coordinates in visual space can be inferred for each voxel in hV5/MT+ using the CF modeling method. Therefore, we can plot the visual field coverage maps that correspond to the CFs of area hV5/MT+ voxels and compare them with the visual field coverage maps obtained during hV5/MT+ pRF mapping (Figure 5D).

#### Statistical Analysis

A two-sample Kolmogorov-Smirnov test was performed in order to compare the pRF size distributions between patients and the AS control subjects. The significance level selected to reject the NULL hypothesis (same distributions) was estimated by comparing the distribution of each AS control subject with the average distribution of all control subjects. The minimum p-value of these comparisons was then used to test for significance between the mean distribution of the AS controls and the patients. We note that this is a conservative choice, and may suppress the identification of small differences.

## Results

### Mapping the pRF boundaries separately for each direction of motion of the visual stimulus

Differences in the retinotopic maps of normal subjects have been observed when the visual stimulus is masked to simulate retinal or cortical scotomas compared to when the full visual field is stimulated (Haak et al., 2012; Binda et al., 2013; Papanikolaou et al., 2015). These biases are important to know in order to ensure that changes in retinotopic organization seen in patients are not simply an artifact of model estimation caused by incomplete stimulus presentation due to the presence of the visual field defect. In previous work we obtained responses in area hV5/MT+ after masking the upper left quadrant of the visual field in healthy subjects, thus simulating an upper left quadrantanopia (artificial scotoma or AS) (Papanikolaou et al., 2015). We found that when the full bar stimulus model is used to estimate the pRFs in these subjects, hV5/MT+ activity extends well within the region of the AS, even though de-facto there was no stimulation there. These erroneous estimates occur for both direct-fit methods (Figure 1A) (Dumoulin and Wandell, 2008) and topography based methods (Figure 1B) (Lee et al., 2013). These biases are not the result of a trivial methodological artifact or eye-movement deviations, but originate from asymmetric BOLD responses occurring when the bar stimulus moves from seeing to non-seeing locations of the visual field (Figure 7 in Papanikolaou et al., 2015) versus vice versa. In Papanikolaou et al. (2015) we argued that the origin of these effects is more likely due to hemodynamical reasons than due to neuropsychological anticipation, based on the fact that they do not occur across visual hemifields.

**Figure 1:**
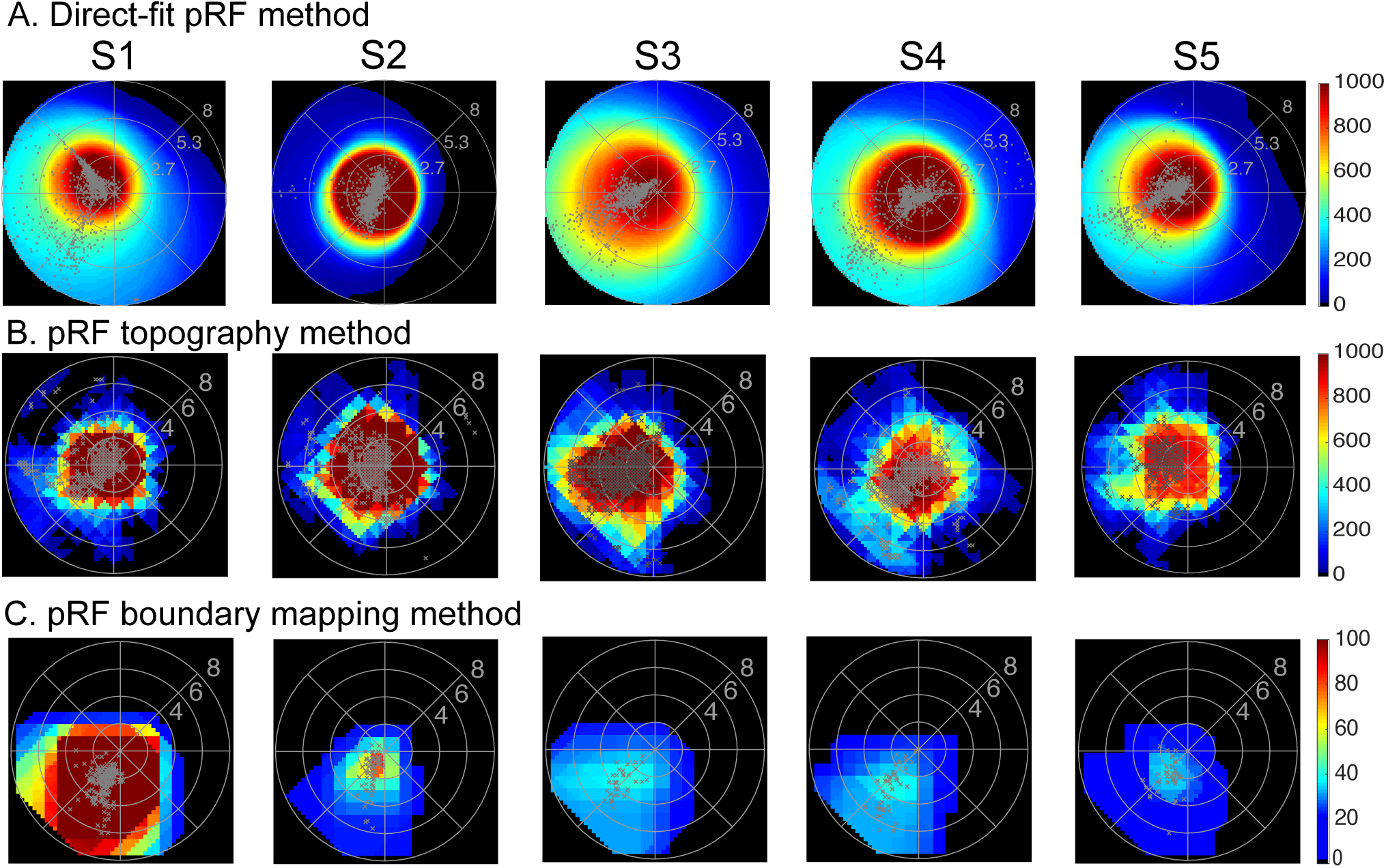
hV5/MT+ visual field coverage maps in Artificial Scotoma subjects. Visual field coverage density maps of area hV5/MT+ of the right hemisphere in all control subjects with an AS at the upper left quadrant using a direct-fit pRF method (**A**), a topography-based method (**B**) and the proposed pRF boundary mapping method (**C**). The color map indicates the number (#) of pRFs that cover each visual field location. The pRF centers from all voxels are plotted as grey dots. The snapshot at the top right corner shows the location of the artificial scotoma in the visual field (grey shaded area). For both direct-fit and topography-based pRF methods, hV5/MT+ maps cover significantly the area of the AS at the upper left quadrant when the full bar stimulus is used for modeling the pRFs. In contrast, for the proposed method (C) in which we map the pRF boundaries directly from the BOLD signal, pRFs in hV5/MT+ are confined to the lower visual field quadrant outside of the AS in all subjects. Activity only modestly crosses the border of the AS (~2 deg) commensurate with the subject’s fixation eye movements. Note that the boundary mapping pRF method is more strict than the direct-fit and the topography-based methods as only well-defined pRFs are retained for analysis. Nevertheless, applying a more strict criterion (variance explained > 30%) for the direct-fit and topography-based method does not eliminate the pRF biases observed within the AS.

In patients, retinotopic mapping is performed using a full bar stimulus (in this case a drifting flickering checkerboard bar), which overlaps the area of the scotoma, and may therefore be prone to similar artifacts. Thus a different approach is needed for comparing responses between patients and AS subjects when using the drifting bar stimulus. We developed a method, which calculates directly the boundaries of the pRF from the BOLD time series of each voxel separately for each direction of motion of the visual stimulus (Methods; Figure S1). In this way, hysteresis phenomena in the BOLD signal that are produced when the bar moves from seeing to non-seeing locations of the visual field can be eliminated. Using this method, pRF estimates in area hV5/MT+ of subjects with a simulated AS, are confined in visual field locations outside of the AS, as expected (Figure 1C).

Subsequently, we used this method for estimating pRF responses in spared area hV5/MT+ of patients with V1+ or optic radiation lesions that resulted in dense contralateral scotomas.

### Patients: Anatomical lesion and visual_field defects

We examined 5 patients, P1–5, with homonymous dense visual field defects within one quadrant of the visual field (visual sensitivity<-20dB). Each patient’s lesion and consequent visual field defect is presented in detail in Table 1. The anatomical location of the lesion and patients’ perimetry tests are presented in Figure 2.

**Figure 2:**
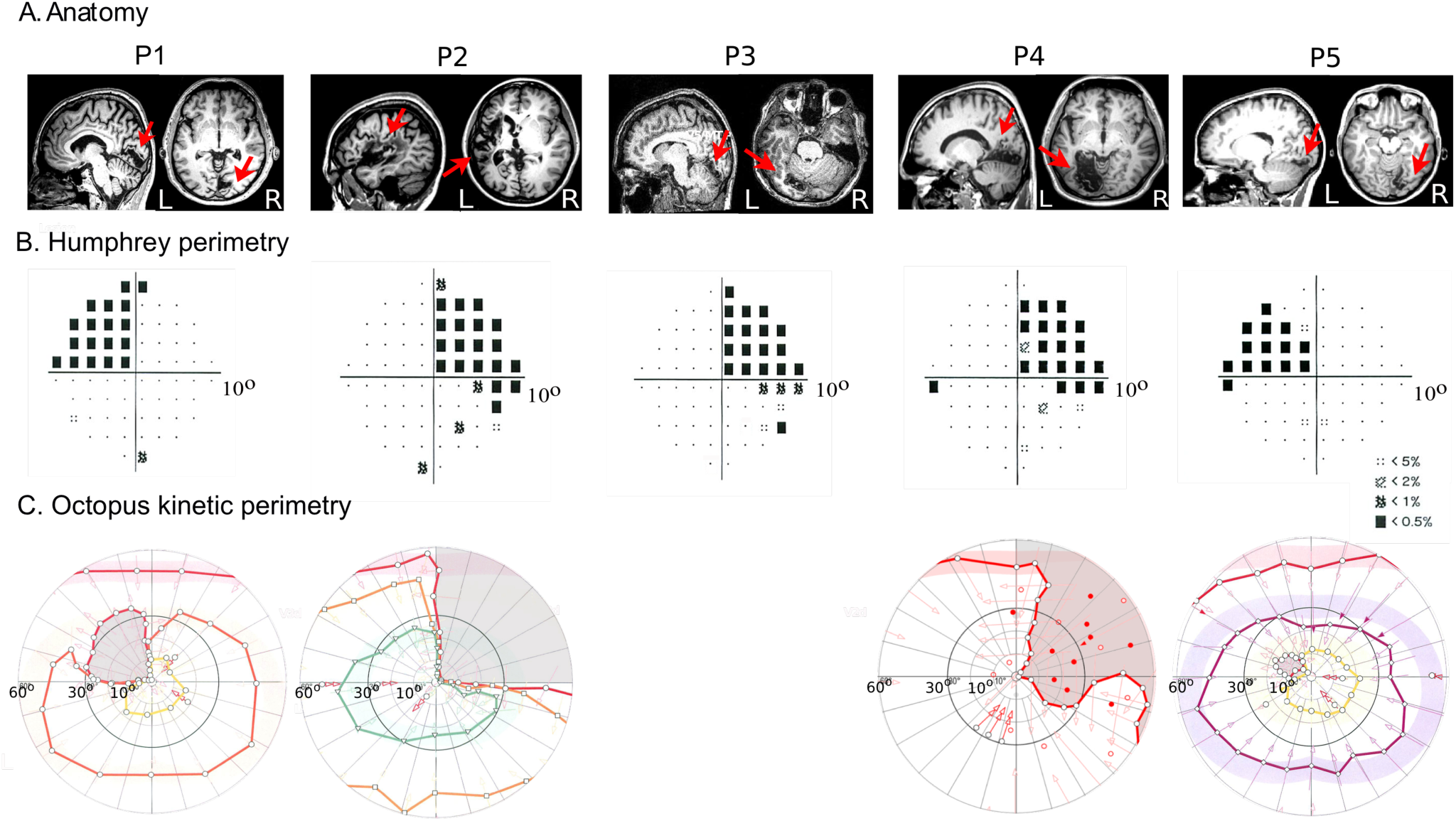
Anatomical location of the lesion and visual field perimetry tests. **A**. Anatomical location of the lesion. A sagittal (top) and an axial (bottom) slice illustrates each patient's anatomical lesion (a red arrow points to the lesion). **B**. Pattern deviation probability plots of the 10-degree Humphrey type (10–2) visual field test for all patients. The small black dots show the locations in the visual field that are normal, while the black squares indicate a visual field defect on a p<0.5% level according to the pattern probability plot (this means that less than 0.5% of normal subjects would be expected to have such a low sensitivity at this visual field location). Pattern deviation numeric plots for all patients had visual sensitivity <-20dB (absolute visual field scotoma) at all visual field locations within the affected quadrants. Black square locations outside the affected quadrants showed visual sensitivity <-10dB (mostly still <-20dB). **C**. Binocular semi-automated 90° kinetic perimetry (Octopus 101; methods) for patients P1, P2, P4 and P5. The area of absolute visual field loss is shaded in light grey.

In brief, patients P1, P3 and P4 have lesions which extend from the part of V1 inferior to the calcarine sulcus to extrastriate ventral visual areas, resulting in contralateral superior quadrantanopic defects (Figure 2A-C.a,c,d). P2 has a superior quadrantanopia (Figure 2B.b) following a temporal optic radiation lesion. P5 has a comparatively smaller lesion, which involves part of the foveal ventral V1 and ventral extrastriate areas V2 and V3 resulting in a dense defect within the left upper visual field quadrant (Figure 2B.e). Patients’ P1, P2, P3 and P4 V1 organization has been described in more detail before (Papanikolaou et al., 2014).

**Table 1: Patient data**. Patient identification (ID), side of brain lesion (Hemisphere), visual areas affected by the lesion (Areas), location of homonymous visual field defect (LUQ: Left Upper Quadrant, RUQ: Right Upper Quadrant) and time span between brain lesion and examination (Δt). P1 has a left superior quadrantanopic defect following a lesion of the right inferior calcarine cortex. The lesion extends from the part of V1 inferior to the calcarine sulcus to extrastriate cortex corresponding to the ventral visual areas V2 and V3. P2 has a circumscribed defect within the right upper visual field quadrant due to/as a consequence of a left-hemispheric infarction of the temporal optic radiation. This results in deafferentiation of a significant portion of V1 by cutting its input, while the gray matter of this area has remained intact. Patient P3 has a homonymous superior quadrantic defect of the right visual field following a lesion of the left V1 inferior to the calcarine sulcus including extrastriate ventral visual areas V2, V3 and V4. P4 has a lesion of the left inferior calcarine cortex, which involves ventral striate area V1, ventral extrastriate areas V2, V3 and V4, and extends to the dorsal area V1 where it spares a small part of the dorsal periphery. This has created a homonymous superior quadrantic defect of the right visual field. P5 has a lesion in the right hemisphere, which involves part of the foveal ventral V1 and ventral extrastriate areas V2 and V3, resulting in a dense defect within the left upper visual field quadrant.

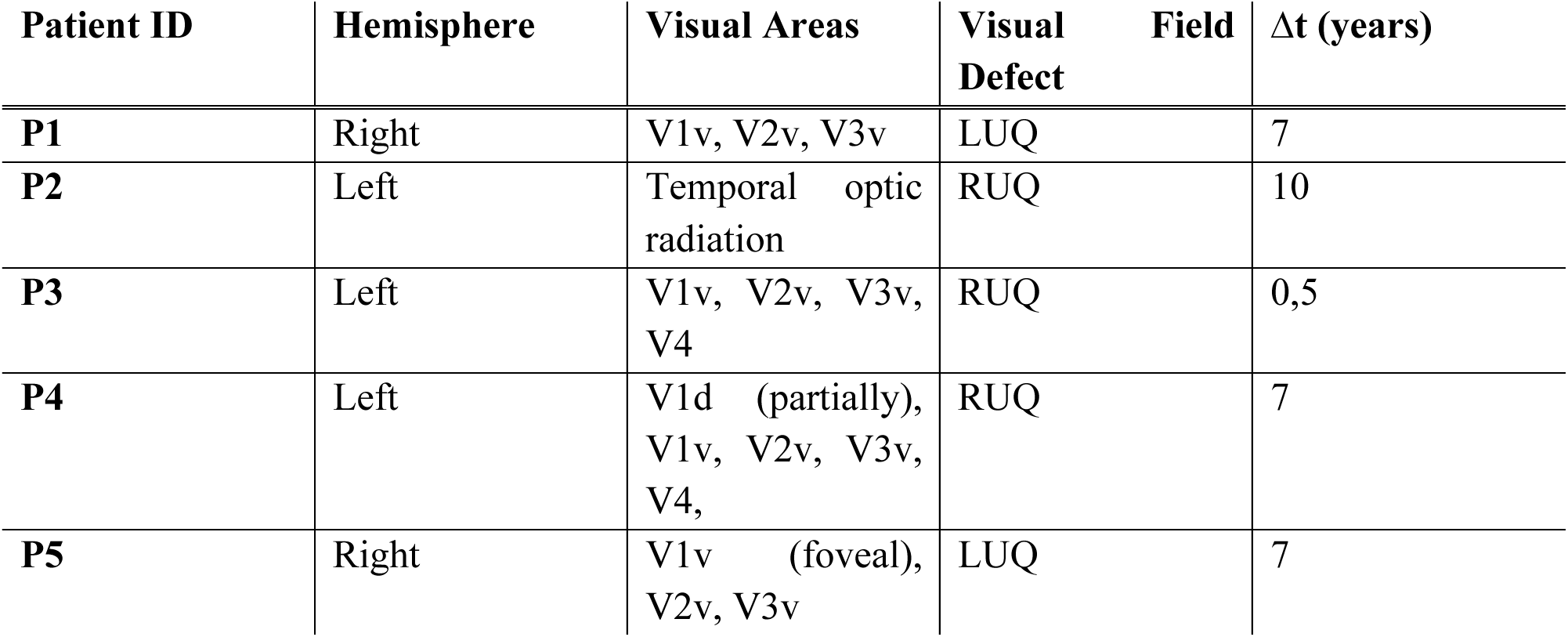

### hV5/MT+ responses following V1+ lesions

We used a direct-fit pRF method (Dumoulin and Wandell, 2008) to identify area hV5/MT+ for all subjects (Figure S2). The identified region of interest was then used to obtain pRF estimates using the proposed mapping method. We compared hV5/MT+ coverage density maps for all patients with the visual perimetry maps defining the perceptual scotoma and with the hV5/MT+ coverage density maps of AS controls. As shown before, pRF estimates in area hV5/MT+ of healthy subjects with a simulated AS at the upper left quadrant, are confined in visual field locations outside of the AS (Figure 1C, Figure 3A.a). In contrast, all patients showed activity in hV5/MT+ that extended well beyond the border of the scotoma into the superior (anopic) visual field quadrant (Figure 3A, red arrows). Raw BOLD responses from voxels in hV5/MT+ confirm that activity arises from stimulus presented within the visual field scotoma (Figure 3B-C). Specifically, we compared the raw BOLD signal change between patients and the AS controls when the stimulus moves from non-seeing (scotoma) to seeing locations of the visual field. In AS controls, the average BOLD signal change drops to baseline values when a horizontal bar is in the superior quadrant (location of the AS; elevation>0, Figure 3B.a). Activity starts when the bar is near 2° from the horizontal meridian (AS border), commensurate with the subject’s fixation eye movements. In contrast, for all patients, activity starts when the stimulus is located well within the perceptual scotoma (elevation>3°, red arrows, Figure 3B.b-f). Moreover, when a vertical bar moves from the ipsilateral to the contralateral visual hemifield, activity starts when the stimulus is located about 2–3° within the ipsilateral visual hemifield, for both patients and AS control subjects. This suggests that, hV5/MT+ activity in the lesioned hemisphere originates from stimulus positions located within the scotoma, and less likely from the contralateral hemisphere.

**Figure 3:**
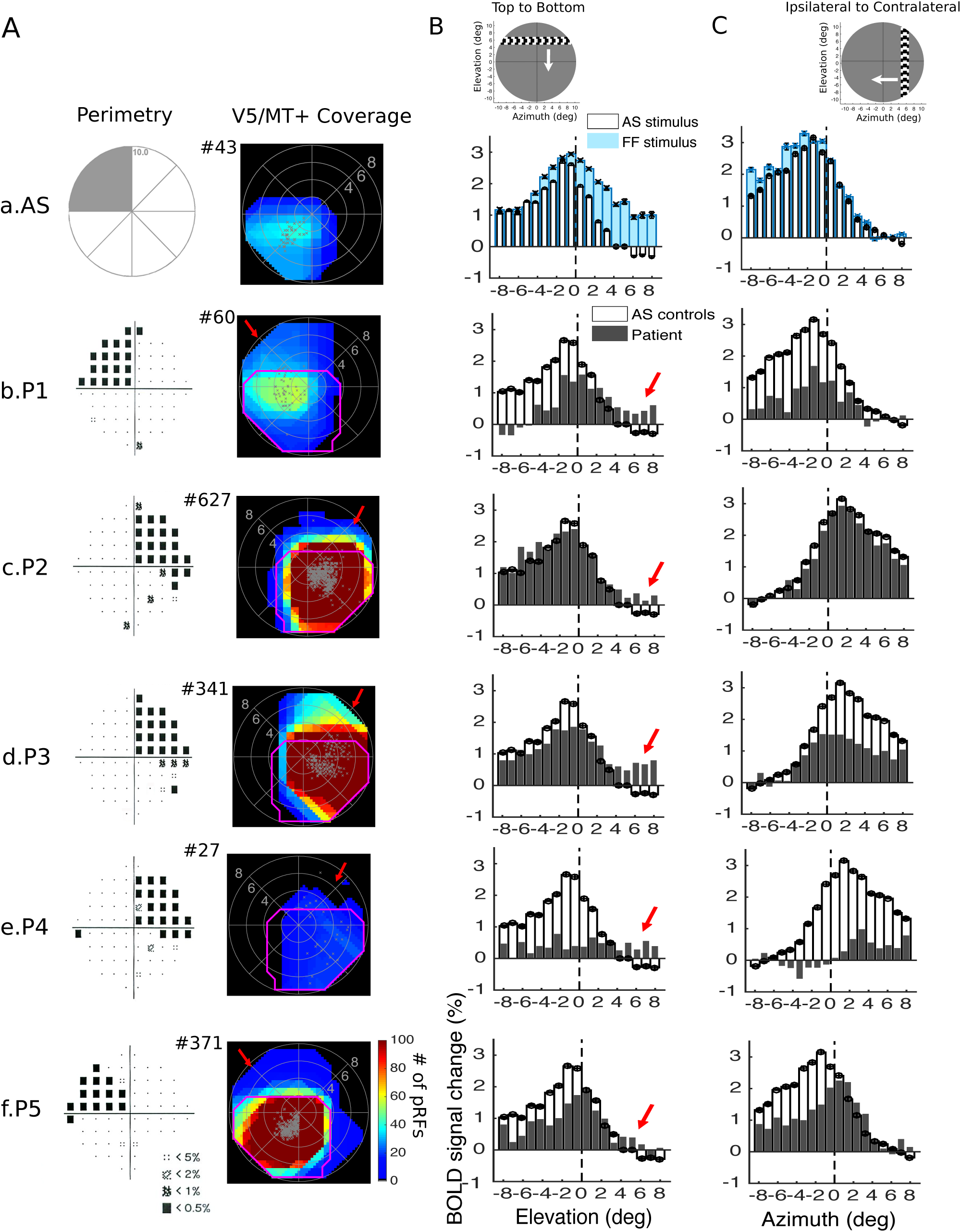
Visual field coverage maps and average BOLD signal change in hV5/MT+ of the lesioned hemisphere. **A**. *Left:* Pattern deviation probability plots of the 10-degree Humphrey type visual field test for all patients as shown in Figure 2B. *Right:* Visual field coverage density maps of area hV5/MT+ of the right hemisphere of an AS control subject (a) and of the lesioned hemisphere for each patient obtained using the proposed pRF boundary mapping method (b-f). To allow comparison between subjects, the scale of the color map has been clipped to the average significantly activated number of voxels of hV5/MT+ of AS controls (97.8+89.15). The total number of significantly activated voxels in hV5/MT+ of each subject is indicated next the graphs with a # symbol. The pRF centers from all voxels within each area are plotted as grey dots. The average coverage density map of all AS controls is overlaid on top of the maps of each patient with magenta color. In contrast to the AS controls, the visual field coverage maps of hV5/MT+ of all patients cover areas that overlap with the dense visual field scotoma (red arrows). B. The average BOLD signal change from all voxels in the right hV5/MT+ in controls and the hV5/MT+ of the lesioned hemisphere in patients as a horizontal bar is moving from the top (elevation>0; AS/scotoma) to the bottom of the visual field (elevation<0; seeing quadrant). Before averaging, the BOLD time series of each voxel is deconvolved to remove the hemodynamic response function (Methods) and the baseline is removed. The baseline here is defined as the signal value when the vertical bar is located in the far ipsilesional part of the visual field, which should produce little or no visual modulation in the region examined. This is calculated as the average BOLD signal change over 5 steps of the bar when the horizontal bar was located between 7–10° in the hemifield ipsilateral to our ROI. This procedure sets the baseline of each voxel to zero. (a) The average signal of the AS controls (white bars) is compared with the full field stimulus condition (blue bars). When the AS is applied, the average BOLD signal change when the bar is in the superior quadrant (location of the AS; elevation>0) drops to baseline values compared with the average signal under the full field stimulus condition. Activity starts when the bar is near 2° from the horizontal meridian (AS border), commensurate with the subject’s fixation eye movements. (b-f) The average signal of the patients (gray bars) compared with the AS controls (white bars). For all patients, activity starts when the stimulus is located well within the perceptual scotoma (elevation>3°, red arrows) in contrast to the AS controls. The error bars indicate the standard error of the mean across control subjects (N=5). The left column shows a snapshot of the orientation of the bar and direction of motion (white arrow). C. Same as in (B), the average BOLD signal change from all voxels in the right hV5/MT+ in controls and the hV5/MT+ of the lesioned hemisphere in patients as a vertical bar is moving from the contralateral (azimuth>0) to the ipsilateral visual hemifield (azimuth<0). For all patients, activity starts when the stimulus is located about 2–3° within the ipsilateral visual hemifield, similar to the AS control subjects. This suggests that, hV5/MT+ activity in the lesioned hemisphere originates from stimulus positions located within the scotoma, and less likely from the contralateral hemisphere.

### Pathways contributing to hV5/MT+ activity following V1+ lesions

To understand the possible source mechanisms of the hV5/MT+ activity observed within the scotoma in patients, we compared hV5/MT+ responses with the responses obtained from area V1. In healthy subjects under full field stimulation, areas V1 and hV5/MT+ cover the contralateral hemifield (Figure 4A-B, top). If the stimulus is masked at the upper left visual field quadrant (Artificial Scotoma or AS), simulating an upper left quadrantanopia, areas V1 and hV5/MT+ cover only the lower visual field quadrant (Figure 4A-B, bottom). Activity only modestly crosses the border of the AS (~2 deg) commensurate with the subject’s fixation eye movements.

**Figure 4:**
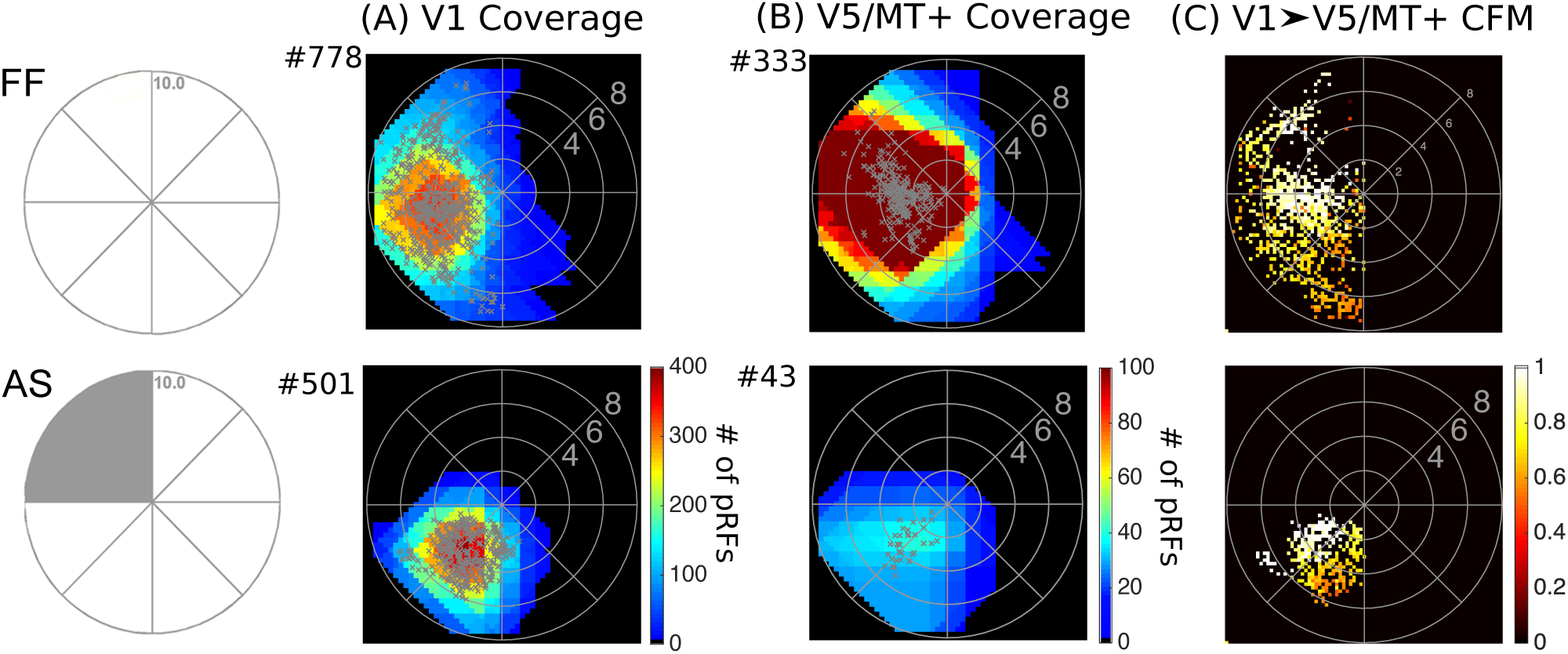
Visual field coverage maps of V1 and hV5/MT+ in Control Subjects. *Top:* Visual field coverage density maps of area V1 (**A**) and the hV5/MT+ complex (**B**) of the right hemisphere of a control subject under Full Field (FF) stimulation. The pRF centers from all voxels within each area are plotted as grey dots. Both areas cover the entire left visual hemifield, as expected. *Bottom:* Visual field coverage density maps of V1 (A) and hV5/MT+ (B) of the right hemisphere of a control subject under Artificial Scotoma (AS) condition. No activity is observed within the area of the AS in control subjects for both V1 and hV5/MT+. **C**. Visual field coverage maps of hV5/MT+ based on the connective field modeling method (CFM) for a control subject under FF stimulation (top) and under the AS condition (bottom). These maps plot, for each voxel in hV5/MT+, the pRFs of the voxels corresponding to the CF center and CF Gaussian spread in the cortical surface of V1. The color map indicates the CF weight so that 1 corresponds to the CF center. CFM coverage maps in control subjects are commensurate to the pRF coverage density maps suggesting that hV5/MT+ activity arises chiefly from area V1.

To further investigate the dependence of hV5/MT+ responses on V1 we used population connective field (CF) modeling. CF modeling is used to predict the activity of voxels in hV5/MT+ as a function of activity in spared V1 (Haak et al., 2013) (Methods). Using a combination of CF modeling and our method for estimating the pRFs, we derive visual field coverage maps for area hV5/MT+ based on the CF location on the cortical surface in V1 (Figure 4C). Specifically, for each voxel in hV5/MT+ we plot the pRF center locations in the visual field of the corresponding CF in V1. We compare the maps derived using the CF modeling with the visual field coverage maps derived using our pRF boundary mapping method. This way, we can estimate which parts in area V1 drive the responses in hV5/MT+.

In AS controls, the coverage maps derived using the CF model cover only part of the lower visual field quadrant, outside of the AS and commensurate with the pRF coverage density maps (Figure 4B-C, bottom). Essentially all functional voxels in hV5/MT+ were linked with a well-defined CF in area V1 at the chosen threshold (explained variance>0.125). This suggests that hV5/MT+ activity in AS controls arises from the visually responsive part of area V1. No activity is observed within the area of the AS. Using this combination of the pRF boundary method and CF modelling, we identified two possible source mechanisms for the hV5/MT+ activity observed with the scotoma of the patients.

### Visual field regions overlapping with the patients ’ scotoma covered by <u>both</u> hV5/MT+ and V1

For patients P2, P3 and P5, most activated visual field locations in hV5/MT+ overlapping with the patients’ perceptual scotoma are also covered by V1 (Figure 5B-C.a,b,c; red arrows). The observed V1 activity within the scotoma is not ectopic but reflects islands of V1 that are spared or only partially damaged (Papanikolaou et al., 2014). It is therefore possible that hV5/MT+ responses corresponding to the scotoma arise from the spared part of area V1. To confirm this, we plotted the hV5/MT+ coverage maps derived using the CF models (Figure 5D). CF coverage maps in these patients are commensurate with the pRF coverage density maps (Figure 5D.a,b,c). Specifically, the maps cover visual field locations overlapping with the scotoma suggesting that, most of hV5/MT+ activity arises from spared or partially injured V1 tissue. Surprisingly however, these patients still have a dense (<-20dB) visual field defect corresponding to these locations (Figure 2B-C).

**Figure 5:**
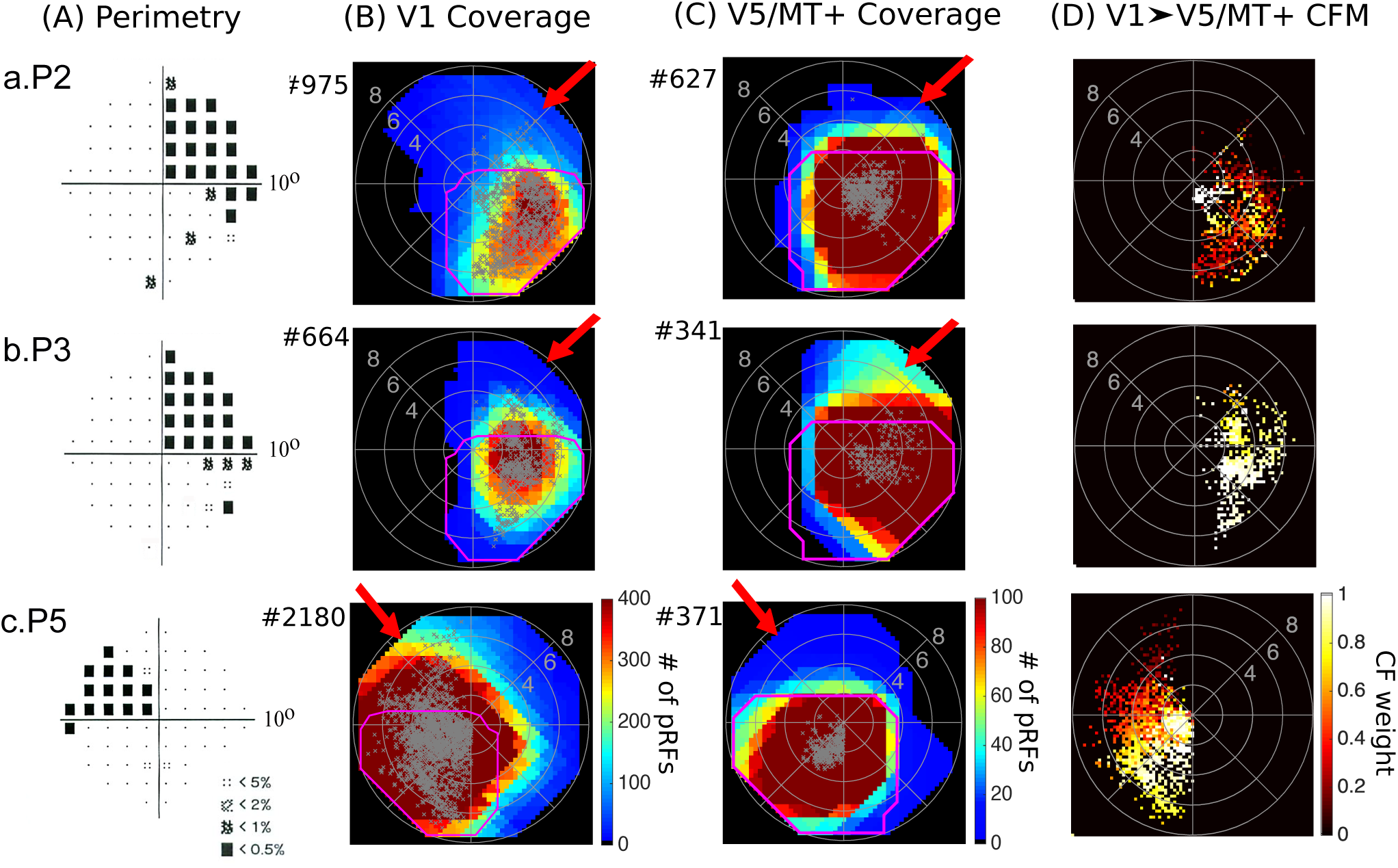
Visual field coverage maps of patients P2, P3 and P5. **A**. Pattern deviation probability plots of the 10-degree Humphrey type visual field test for patients P2, P3 and P5 as shown in Figure 2B. **B**. Visual field coverage density maps of the spared part of area V1 and **C**. the hV5/MT+ complex of the lesioned hemisphere for patients P2, P3 and P5. The scale of the color map has been clipped to the average significantly activated number of voxels plus one standard deviation for V1 (182.4+182.3) and the average significantly activated number of voxels for hV5/MT+ (97.8+89.15) of AS controls. The total number of significantly activated voxels for each subject is indicated next the graphs with a # symbol. The pRF centers from all voxels within each area are plotted as grey dots. The average coverage density map of all AS controls is overlaid on top of the maps of each patient with magenta color. For these patients activity extending beyond the border of the scotoma in hV5/MT+ is also present in V1 (red arrows). Apparently, this activity is not sufficient to mediate conscious vision suggesting it is either too disorganized to elicit a percept or that damage to other areas is responsible for the visual deficit. **D**. Visual field coverage maps of hV5/MT+ based on the connective field modeling method (CFM). CF coverage maps in patients P2, P3 and P5 cover visual field locations overlapping with the scotoma confirming that responses in hV5/MT+ within the scotoma arise from the spared part of area V1.

In principle, the lack of a percept may happen because: i) retinotopically corresponding extrastriate areas are injured, or ii) the activity generated may be too weak, too asynchronous, or too disorganized to elicit a percept. For patients P3 and P5 the lesion includes extrastriate visual areas V2v and V3v. Therefore, the visual field deficit of these patients may be due to damage within these extrastriate areas. Nevertheless, visual information reaches area hV5/MT+ suggesting that the V1 to hV5/MT+ projection is partially spared.

Patient P2, however, has an isolated/exclusive lesion of the optic radiation and thus the pathways from V1 to higher visual areas have survived the lesion. One possibility is that the lesion has affected the projections from V1 to extrastriate visual areas. However, this is unlikely as ventral areas V2 and V3 also showed functional imaging responses that overlap with this patient’s scotoma (Figrue S3), supporting the viewpoint that the extrastriate cortex remains functional in this subject. Therefore, the visual field deficit of this patient cannot be related to damage within these extrastriate areas. In this case, option (ii) may dominate. We previously found (Papanikolaou et al., 2014), that the mean amplitude of area V1 pRF centers that fall inside the scotoma is significantly lower (0.77±0.09; mean±standard deviation) than the mean amplitude of pRF centers that fall in the inferior (seeing) quadrant for this patient (1.06±0.05). The same holds in hV5/MT+ (mean pRF amplitude within the scotoma is 0.83±0.02 compared with 0.92±0.05 in the inferior/seeing quadrant). Note however, that the observed decrease is modest in both areas V1 and hV5/MT+. This suggests that although it is possible that the decrease in the level of visually driven activity may contribute to the loss of visual perception, it is unlikely by itself to be the sole explanation (see below).

### Visual field areas overlapping with the patients ’ scotoma covered by hV5/MT+ but not V1

The visual field coverage density maps of hV5/MT+ in patients P1 and P4, overlap with parts of the visual field scotoma that are not covered by area V1 (Figure 6B-C.a,b; green arrows). In fact, the retinotopically corresponding part of area V1 is anatomically lesioned. This suggests that hV5/MT+ activity within the scotoma in these patients arises via pathways that bypass area V1, i.e. through the SC and pulvinar (Rodman et al., 1990; Weiskrantz, 2004; Barleben et al., 2015) and/or through the LGN (Maunsell et al., 1990; Sincich et al., 2004; Schmid et al., 2009; Schmid et al., 2010). Although these patients have a relatively lower number of pRFs activated in hV5/MT+ than AS controls, activity extends well beyond the border of the scotoma and beyond the activity observed in AS controls. Note that our pRF mapping method is quite conservative and only well-defined pRFs that are above the noise level are mapped. An example of the BOLD activity of a pRF that lies well within the scotoma of these patients is presented in Figure S4.

**Figure 6:**
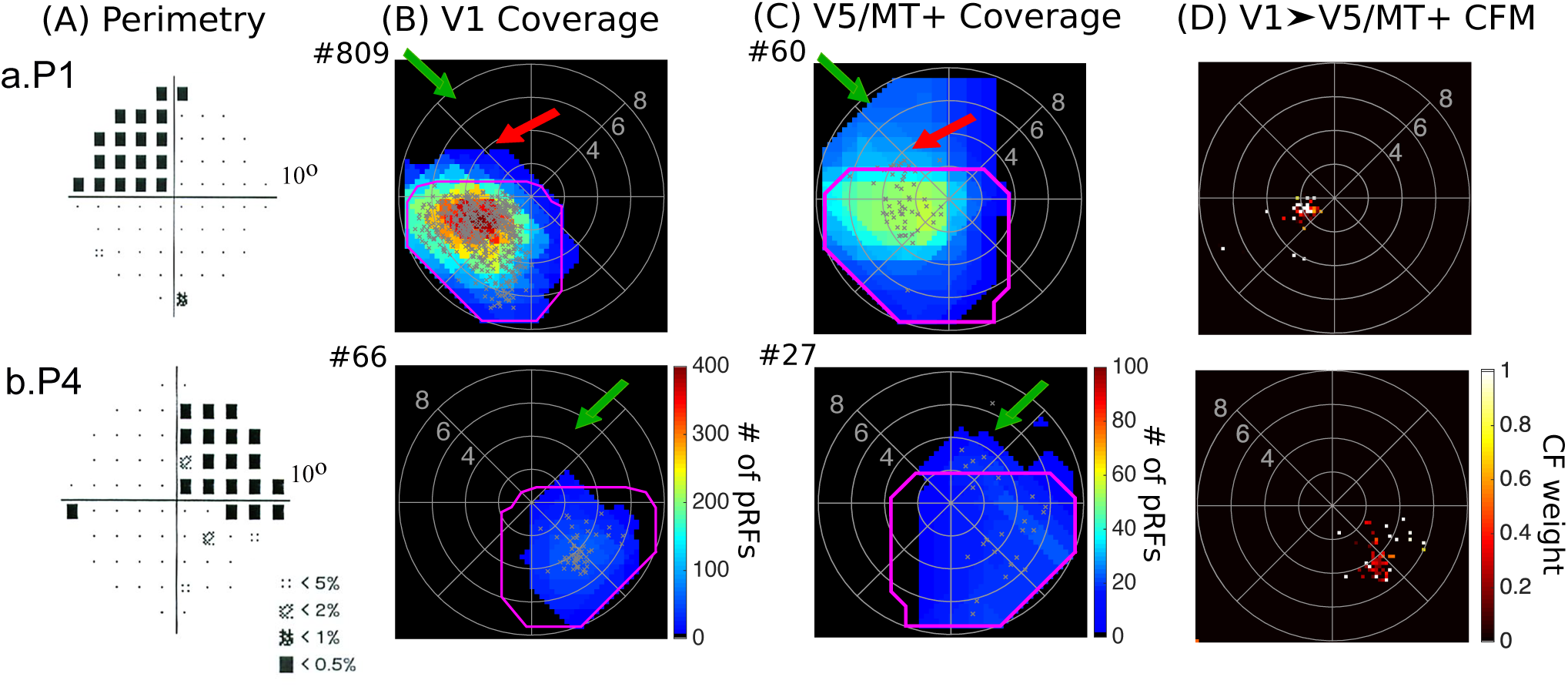
Visual field coverage maps of patients P1 and P4. **A**. Pattern deviation probability plots of the 10-degree Humphrey type visual field test for patients P1 and P4. **B**. Visual field coverage density maps of the spared part of area V1 and **C**. the hV5/MT+ complex of the lesioned hemisphere for patients P1 and P4. The pRF centers from all voxels within each area are plotted as grey dots. The average coverage density map of all AS controls is overlaid on top of the maps of each patient with magenta color. The total number of significantly activated voxels for each subject is indicated next the graphs with a # symbol. Patients P1 and P4 have visual field areas overlapping with the patients’ scotoma that are covered by V5/MT+ but not V1 (green arrows) suggesting the existence of functional V1-bypassing pathways. Activity in hV5/MT+ alone is not sufficient to elicit a percept. Patient P1 also has a part of the visual field overlapping with the patient’s scotoma that is covered by both areas V1 and hV5/MT+ similar to patients P2, P3 and P5. **D**. Visual field coverage maps of hV5/MT+ based on the connective field modeling method (CFM). For patients P1 and P4, the CF modeling links the voxels in hV5/MT+ to voxels in V1 with pRF centers in the inferior (seeing) quadrant only suggesting that responses observed within the scotoma in hV5/MT+ using the pRF mapping method are independent of V1 input.

To confirm that hV5/MT+ activity in these patients does not depend on V1 input, we applied the CF modeling (Figure 6D). A large fraction of visually responsive voxels (13 out of 27 voxels) in hV5/MT+ of patient P4 were not linked with any voxel in V1 using the CF modeling method (variance explained<0.125). These voxels have well-defined pRFs using our pRF mapping method but no connective field in V1, confirming that they receive their inputs from V1 by-passing pathways. However, input from these pathways is apparently not sufficient to mediate conscious vision. Indeed, the mean amplitude of hV5/MT+ pRFs with center within the scotoma of patient P4 is smaller than the mean hV5/MT+ pRF amplitude in the healthy hemisphere (ratio: 0.67±0.10). However, weak modulation may contribute to the loss of vision, but cannot explain it: the inferior (seeing) quadrant outside the visual field scotoma has pRFs with a similar mean amplitude (ratio: 0.68±0.10), without an obvious visual field deficit. It is also evident that in normal subjects a modest decrease in the contrast of the stimulus may induce a similar drop in the level of activity in area hV5/MT+ or other extrastriate areas without compromising visual perception. A reasonable hypothesis is then that either i) activity in area V1 is itself important for visual perception, or ii) in the absence of V1 activation the activity elicited in the extrastriate cortex (here hV5/MT+) is not appropriately organized/synchronized to support visual perception.

It is worth mentioning here that patient P4 appears to have a small amount of spared vision across the vertical meridian, as shown in the Octopus perimetry map (Figure 2C.d). The part of area V1 that corresponds to the vertical meridian in this patient is anatomically lesioned. It is therefore possible that activity observed in area hV5/MT+ at the vertical meridian may contribute to some degree to visual awareness, though other mechanisms are also likely to be involved (e.g. through the contralesional hemisphere).

Similar to patient P4, the hV5/MT+ coverage map of patient P1 covers visual field locations overlapping with the patient’s scotoma that are not covered in V1 (Figure 6B-C.a, green arrows). However, this patient also shows locations of the visual field that are covered by both areas V1 and hV5/MT+, similar to patients P2, P3 and P5 (Figure 6B-C.a, red arrows). CF modeling in this patient suggests that hV5/MT+ activity may arise from a mixture of input mechanisms (Figure 6D.a). This is described in detail in the next section.

It is important to note that the overlap between visual field coverage maps and the scotoma seen on perimetry cannot be explained by eye movements. Subjects were able to maintain fixation within 1.5° radius from the center of fixation except for very occasional excursions beyond this range (Figure S5). The results remain unchanged after removing from the analysis these epochs, where the subjects had eye deviations (>1.5°) from the fixation point. Patient’s P3 eye movements were not recorded, and therefore we cannot completely exclude the possibility that activity observed within the scotoma of this patient is the result of eye movements. However, several facts oppose this hypothesis. First, results observed in this subject were in line with the observations made in patients P1, P2 and P5 that have undergone rigorous eye movement tracking. Second, the subject performed a challenging detection task at fixation and his performance was maintained >80% correct. Finally, the retinotopic maps of his healthy hemisphere were well-organized suggesting that he did not make frequent large, confounding, eye movements that could explain the large (>1.5°) hV5/MT+ pRF coverage within the scotoma.

### Cortico-cortical interactions between hV5/MT + and spared V1

To better understand the source of activation within the scotoma in hV5/MT+ of patients, we plot the visual field elevation (y coordinate) of the hV5/MT+ pRF centers against the pRF elevation of the V1 voxels corresponding to the hV5/MT+ voxels' CF centers (Figure 7). A testament that the CF modeling is an appropriate method for measuring the dependence between hV5/MT+ and V1 responses, is that the visual field location estimates in hV5/MT+ of AS controls as measured by the CF method are highly correlated to the location estimates derived using the pRF method (Figrue S6, correlation coefficient r=0.59±0.07, p=10^−4^). This result is expected if hV5/MT+ voxels receive their dominant input from V1.

**Figure 7:**
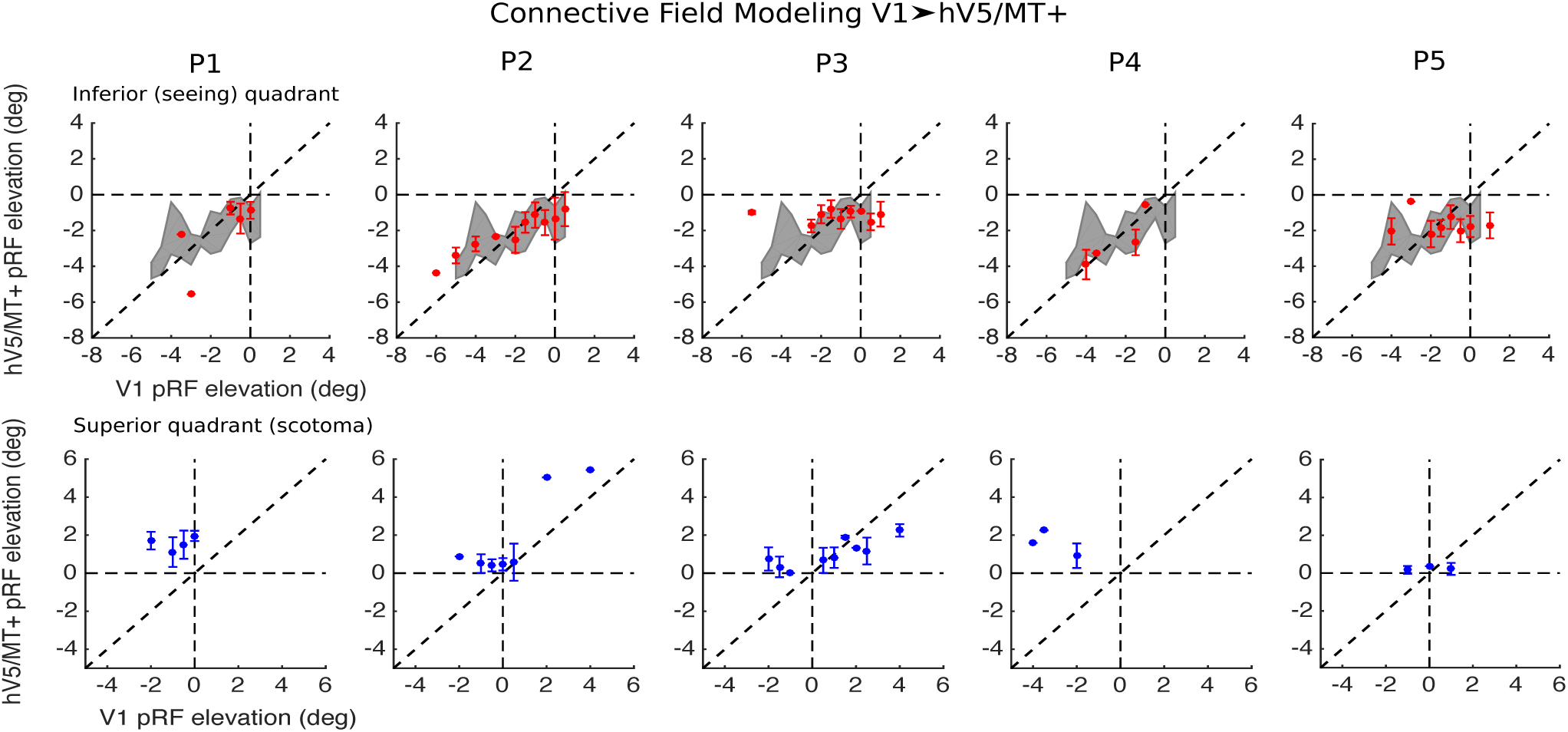
Cortico-cortical connectivity between hV5/MT+ and V1 in patients. *Top row, red:* The location of voxels in hV5/MT+ with pRF center elevation in the inferior (seeing) quadrant (y<0) is plotted as a function of the pRF center elevation of the corresponding CF center in V1 (as found using the CF modeling method, see methods). The grey shaded area represents the mean±standard deviation of the AS controls (N = 5). *Bottom row, blue:* The location of voxels in hV5/MT+ with pRF center elevation in the superior quadrant (scotoma, y>0) is plotted in blue as a function of the pRF center elevation of the corresponding CF center in V1. The error bars indicate the standard deviation across voxels within an elevation bin (bin size = 0.5°) for each patient. For patients P1 and P4, voxels in hV5/MT+ that have pRF centers within the scotoma (y>0; superior quadrant) are linked (in the connective field sense) *only* with voxels in V1 that have pRF centers in the inferior (seeing) quadrant. This correspondence is retinotopically ectopic confirming that the retinotopically corresponding V1 voxels have been lesioned. This ectopic association suggests that visually driven hV5/MT+ responses within the scotoma do not have their source in visual responses of surviving V1 voxels, but instead arise through V1-bypassing pathways. Note that in the control subjects with the artificial scotoma this situation does not arise. For patients P2 and P3, voxels in hV5/MT+ that have pRF centers within the scotoma (y>0) are linked with voxels in V1 whose pRF centers belong to either the superior (scotoma) quadrant or the inferior (seeing) quadrant, suggesting that hV5/MT+ responses within the scotoma may arise within the spared V1 cortex, through V1-bypassing pathways, or via a combination of both.

As described before, CF coverage maps for patients P2, P3 and P5, cover visual field locations overlapping with the scotoma suggesting that, most of hV5/MT+ activity arises from spared or partially injured V1 tissue (Figure 5D). A significant correlation between the location estimates derived by the pRF and the CF method is observed for hV5/MT+ pRFs that fall within the scotoma for patients P2 and P3 (P2, r=0.64 and P3, r=0.52, p=10^−2^; Figure 7, bottom row) confirming that hV5/MT+ responses in these patients arise, at least in part, from the spared V1. Note that essentially all functional voxels in hV5/MT+ were linked with a well-defined CF in area V1 at the chosen threshold (explained variance>0.125). However, there were also some hV5/MT+ voxels with pRF centers inside the scotoma that were linked to V1 voxels with pRFs in the inferior (seeing) quadrant (Figure 7, bottom row). The disparity between visual field location as predicted by the CF modeling and pRF mapping suggests that there are additional pathways contributing to hV5/MT+ activity. It is therefore possible that a small fraction of hV5/MT+ voxels in these patients receive their input from V1-bypassing pathways, but this hypothesis cannot be fully confirmed here. Patient P5 has only a few voxels (7) with pRF centers falling within the scotoma area near the horizontal meridian, which is not enough to assess whether there is a significant correlation in the location estimates between the two methods (Figure 7, bottom row). However, all voxels in area hV5/MT+ of this patient had a well-defined CF suggesting that, hV5/MT+ activity in this patient arises from the spared V1.

On the other hand, for patients P1 and P4, CF modeling yields a more complex picture. A large fraction of visually responsive voxels (13 out of 27 voxels) in hV5/MT+ of patient P4 were not linked with any voxel in V1 using the CF modeling method (variance explained<0.125). These voxels have well-defined pRFs using our pRF mapping method but no connective field in V1, suggesting that they receive their inputs from V1 by-passing pathways. Interestingly, the remaining hV5/MT+ voxels that have a well-defined CF are linked to voxels in V1 with pRF centers that lie exclusively in the inferior (seeing) quadrant (Figure 6D.b). This holds true even for hV5/MT+ voxels with pRF centers that lie well within the superior quadrant (i.e. >2° inside the scotoma, Figure 7, bottom row). Since the pRF size of V1 voxels is relatively small (1–2°), pRFs that lie in the inferior quadrant do not extend more than 1–2° within the superior quadrant. This suggests that responses observed within the scotoma (>2° from the horizontal meridian) in hV5/MT+ of this patient using the pRF mapping method cannot be explained by surviving V1 input.

For patient P1, most visually responsive voxels in hV5/MT+ have a well-defined CF. The CF method links the voxels in hV5/MT+ that have pRFs inside the scotoma to voxels in V1 that have pRFs in the inferior (seeing) quadrant (Figure 7, bottom row). This suggests that hV5/MT+ voxels do receive part of their input *ectopically* from spared V1 voxels. However, hV5/MT+ voxels with pRFs centers inside the scotoma also show responses when the visual stimulus is well inside the blind quadrant (see Figure 3B), which cannot arise from the ectopic V1 locations, which respond to stimuli in the inferior (seeing) quadrant. In fact, the strongest responses (the pRF peak) of hV5/MT+ voxels arise within the scotoma, whereas from their connective fields we would conclude it should lie in the inferior (seeing) quadrant. This argues that responses within the scotoma in these hV5/MT+ voxels arise chiefly from V1-bypassing pathways.

For the inferior (seeing) quadrant, the CF method yields similar hV5/MT+ location estimates to the pRF mapping method. In particular, the location estimates derived by both methods were found to be within the range of the AS controls for all patients (Figure 7, top row). Moreover, a significant correlation is observed for hV5/MT+ pRF centers in the inferior (seeing) quadrant of patients P1, P2 and P4 (r=0.52, r=0.23 and r=0.67 respectively, p<10^−2^, Figure 7, top row), but not for patients P3 and P5 (P3: r=-0.01, p=0.87, P5: r =-0.03, p=0.6). The weak correlation between the CF and pRF estimates in patients P3 and P5 may be because some voxels in hV5/MT+ of the inferior quadrant are linked with V1 voxels that fall within the scotoma, close to the horizontal meridian (Figure 7, top row). These are not necessarily ectopic connections but may emerge from the fact that hV5/MT+ pRFs are large and cover both quadrants near the horizontal meridian. Thus for some voxels, hV5/MT+ activity may partly arise from visually-responsive, spared, V1 regions in the superior quadrant (scotoma) of these patients, an effect that is not feasible in AS-controls, for whom there is no visually driven activity (i.e. no stimulus presented) in the superior quadrant.

In summary, we have identified responses in hV5/MT+ that cover the patients’ scotoma. For some patients such hV5/MT+ responses appear to be mediated by spared V1 to hV5/MT+ projections while, for others, by V1-bypassing pathways or both. Unfortunately, the presence of visually driven BOLD activity in hV5/MT+ is not sufficient to conclude that there is conscious vision. Perhaps more surprisingly, even the presence of visually driven BOLD activity in a (partially lesioned) area V1 region and its retinotopically corresponding early extrastriate areas, including area hV5/MT+, does not guarantee visual perception in some subjects. This suggests that BOLD response activity across the early visual cortex is not a sufficient criterion for deciding whether the capacity for visual perception is spared in subjects with post-geniculate lesions. The fine coordination of activity patterns across visual areas at higher temporal resolution may be an important determinant of whether visual perception persists following lesions of the visual system.

### Population receptive field size in hV5/MT+

We found that the organization of hV5/MT+ in our patients differs from that of AS controls. This does not necessarily reflect reorganization but may be the result of different input sources that are uncovered following V1 lesions. Additionally, we examined whether the pRF size of hV5/MT+ changes in patients with post-geniculate lesions. We found a larger mean pRF size in three out of five patients (P1, P3 and P5; Figure 8A) compared with the AS controls. The effect was significant for patients P3 (p=10^−13^ < p=10^−11^, Kolmogorov-Smirnov test, significance is reported as p = a < b, where b is the value selected to reject the NULL hypothesis; Methods) and P5 (p=10^−12^ < p=10^−11^), but not for patient P1 (p=10^−4^ > p=10^−11^). No significant difference was observed for the remaining patients (P2: p=10^−2^ > p=10^−11^, P4: p=0.11 > p=10^−11^). Significantly larger pRF sizes were observed before near the scotoma border in area V1 of patients P1, P2, P3 and P4 (Papanikolaou et al., 2014). However, in hV5/MT+ the effect is not restricted to pRFs near or inside the scotoma but occurs for pRFs in the inferior (seeing) quadrant as well (Figure 8B), suggesting that area hV5/MT+ might be able to undergo a larger extend of reorganization compared to V1, at least for some patients.

**Figure 8:**
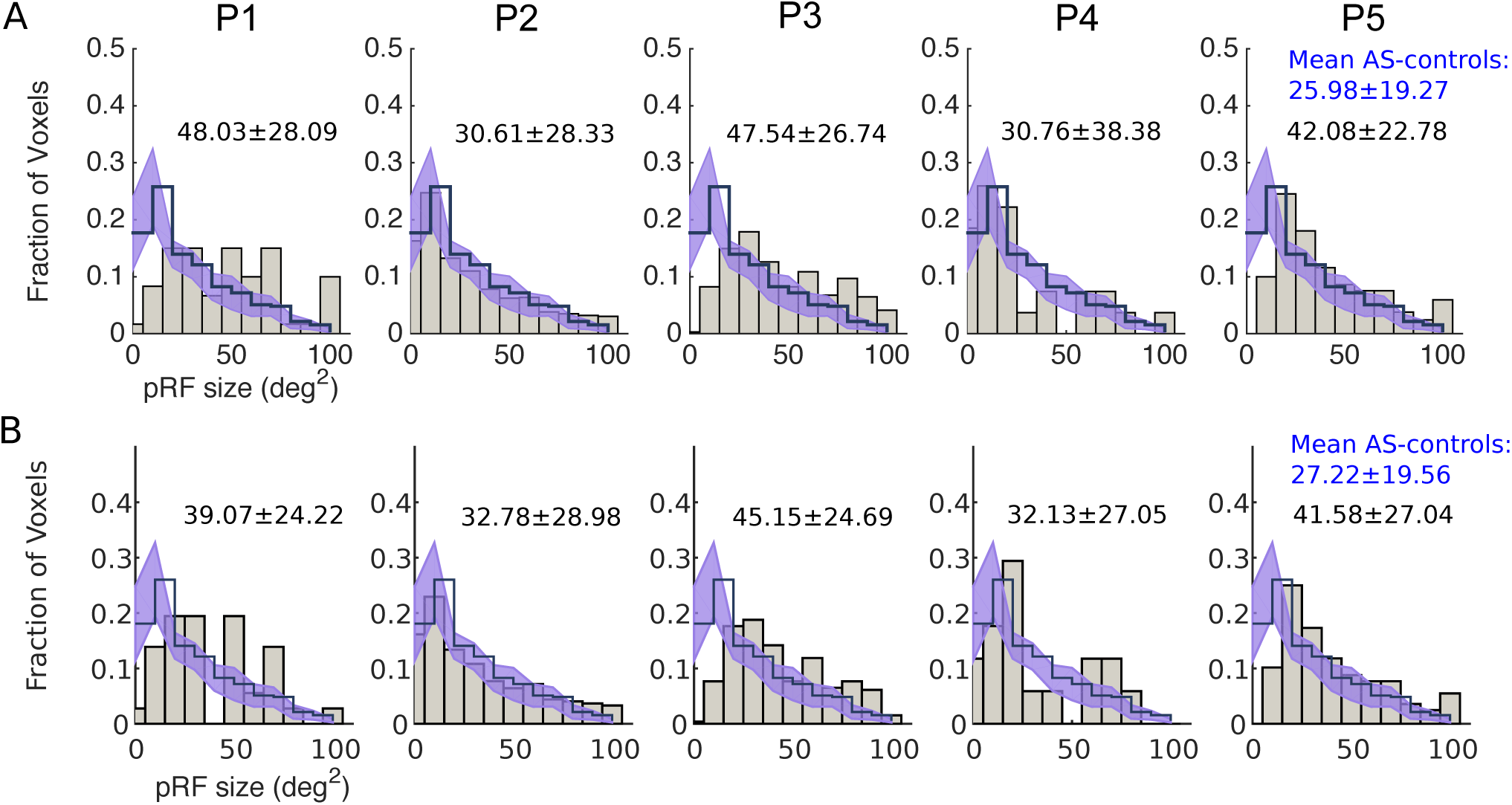
Population receptive field size in hV5/MT+ of the lesioned hemisphere. **A**. Histograms of the distribution of pRF size from hV5/MT+ of all patients (gray bars) compared with the mean distribution of AS controls (step histogram). The shaded area indicates the SEM across the AS controls. The pRF size distribution of patients P1, P3 and P5 is shifted toward larger pRF sizes compared with the AS controls. The mean and standard deviation of each distribution for each patient is indicated on top of the graphs. The mean and standard deviation of the average distribution of AS controls is indicated in blue colour. **B**. Same as in (A) but for pRF centers that are located in the inferior (seeing) quadrant only (pRF elevation < 0).

### hV5/MT+ responses of the contra-lesional hemisphere

Reorganization might occur as well in the contra-lesional, healthy hemisphere, following V1 injury. Previous studies in patients with visual cortical lesions have suggested that residual vision in the blind hemifield might be mediated by visual areas in the intact hemisphere (Ptito et al., 1999; Henriksson et al., 2007; Raninen et al., 2007; Reitsma et al., 2013). Enhanced ipsilateral activation of hV5/MT+ has been observed before in a blindsight patient with extensive V1 injury (Goebel et al., 2001; Morland et al., 2004), and strengthening of callosal connections between the two hV5/MT+ complexes has also been reported (Silvanto et al., 2007; Bridge et al., 2008). We compared the hV5/MT+ pRF coverage density maps of the contra-lesional (healthy) hemisphere for all patients with that of AS controls. In AS controls, the hV5/MT+ complex of the hemisphere ipsilateral to the AS covers one hemifield of the visual field, as expected. Patients P2, P3, and P4 have a significantly smaller hV5/MT+ representation compared with AS controls (AS: # of voxels = 218±104, P2: # of voxels = 67, p = 0.0316, P3: # of voxels = 41, p = 0.0192, P4: # of voxels = 26, p = 0.0147, t-test), suggesting that the pathways to hV5/MT+ of the healthy hemisphere are also influenced by the lesion. Moreover, the pRF coverage density maps of patients P2 and P4 fail to cover the entire visual field hemifield (Figure 9A). Patient P5 on the other hand, shows a significantly enhanced hV5/MT+ representation compared with AS controls (# of voxels = 449, p = 0.008, t-test). The pRF coverage density map of this patient also extended bilaterally, showing increased ipsilateral coverage compared to AS controls (Figure 9A). As each patient has a unique lesion, it is not surprising that different patients show different patterns of activation. The factors that contribute to those differences however, are not entirely clear and more studies with a larger number of patients will be needed in the future to improve understanding of visual processing in the context of injury.

**Figure 9:**
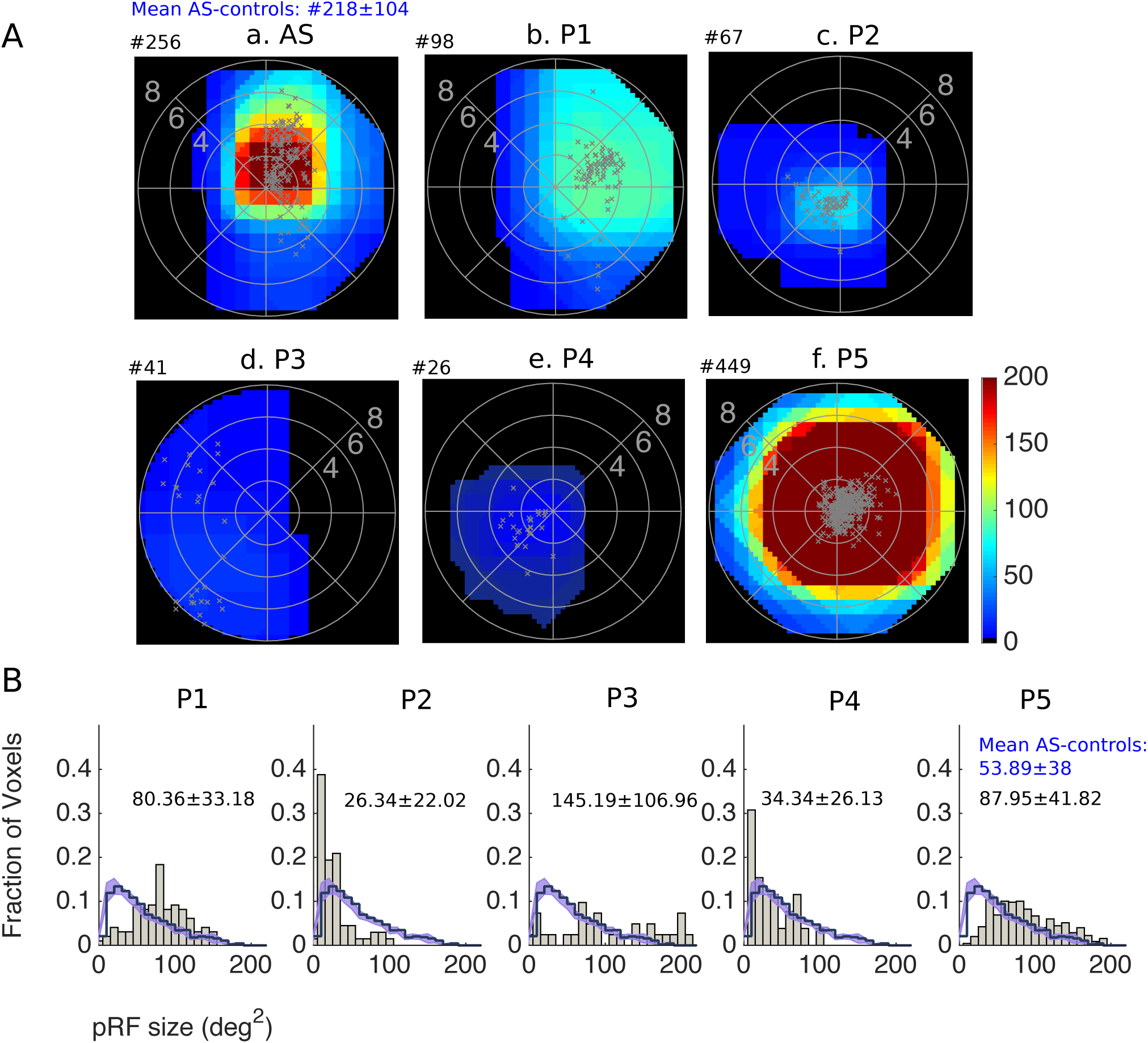
Visual field coverage maps and population receptive field size in hV5/MT+ of the contra-lesional hemisphere. **A**. Visual field coverage density maps of hV5/MT+ of the contra-lesional hemisphere for all patients. The scale of the color map has been clipped to the average significantly activated number of voxels for hV5/MT+ (218+104) of AS controls. The total number of significantly activated voxels for each subject is indicated on top of the graphs with a # symbol. The pRF centers from all voxels within each area are plotted as grey dots. **B**. Histograms of the distribution of pRF size from hV5/MT+ of the contra-lesional hemisphere of all patients (gray bars) compared with the mean distribution of the hemisphere ipsilateral to the AS (left) of AS controls (step histogram). The shaded area indicates the SEM across the AS controls. The mean and standard deviation of each distribution for each patient is indicated on top of the graphs. The mean and standard deviation of the average distribution of AS controls is indicated in blue colour.

Interestingly, the distribution of contra-lesional pRF sizes were different between patients and AS controls (Figure 9B). Specifically, patients P1, P3 and P5 had significantly larger mean pRF size compared with AS controls (Figure 9B, P1: p=10^−16^ < p=10^−06^, P3: p=10^−08^ < p=10^−06^, P5: p=10^−42^ < p=10^−06^, Kolmogorov-Smirnov test). On the other hand, patient P2 had significantly smaller mean pRF size (Figure 9B, P1: p=10^−11^ < p=10^−06^). No significant difference was observed for patient P4 (p = 0.0927 > p=10^−06^). These results suggest that hV5/MT+ of the healthy hemisphere may also undergo significant reorganization following V1 injury, with potential implications in visual rehabilitation.

## Discussion

We measured area hV5/MT+ responses in five patients (P1–5) with chronic post-geniculate lesions resulting in dense homonymous visual field defects.

One important question is whether there are responses in hV5/MT+ to stimuli presented within the scotoma and what pathways these responses arise from. For all patients, visual field coverage density maps of hV5/MT+ overlap with areas of the perimetric scotoma (Figure 3). V5/MT activity after V1 lesions has been observed before in monkeys (Bruce et al., 1986; Rodman et al., 1989; Girard et al., 1992; Rosa et al., 2000; Schmid et al., 2010) and humans (Barbur et al., 1993; ffytche et al., 1996; Schoenfeld et al., 2002; Morland et al., 2004; Bridge et al., 2010; Whitwell et al., 2011). We identified two mechanisms that can account for this activity.

### Responses arising from the spared part of area V1

For 4/5 patients (P1–3, P5) there were visual field regions overlapping with the patients’ perceptual scotoma that were covered by both area hV5/MT+ and area V1. The elicited hV5/MT+ activity, corresponding to these visual field scotoma regions likely arises from the spared part of area V1 (Figure 5). In 3/5 patients (P2, P3, P5), cortico-cortical connective field modeling confirms that voxels in hV5/MT+ that correspond to the visual field scotoma receive inputs from voxels in the spared part of area V1 that also correspond to the visual field scotoma. For the remaining patient (P1), the visual field coverage map of area hV5/MT+ splits into two sections: i) a part that has a corresponding region in the visual field coverage map of area V1 (Figure 5b, red arrows), and ii) a part that does not (Figure 5b, green arrows). For part i) although both V1 and hV5/MT+ areas cover visual field regions overlapping with the patient’s perceptual scotoma, cortico-cortical connective field modeling suggests that hV5/MT+ activity in this region arises, at least in part, by V1-bypassing pathways. For part ii) activity in hV5/MT+ is not associated with activity observed in V1 suggesting that V1-bypassing pathways are dominant in this region.

We showed previously that pRF maps in spared V1 may overlap significantly with dense regions of the perimetric scotoma without contributing to visual awareness (Papanikolaou et al., 2014), and postulated that lesions downstream of V1 input may be responsible. Here we show that even though pRF maps of both area V1 and hV5/MT+ cover the same region of the scotoma, this does not guarantee that visual awareness will be present there (Figure 5). Patients P3 and P5 have a cortical lesion that involves ventral areas V2 and V3 suggesting that the visual field deficit in the superior quadrant of these patients may be due to loss of activity in these areas. This however cannot explain the visual field deficit of patient P2, who has an optic radiation lesion that spares projection pathways of area V1 as well as extrastriate areas. The coverage maps of P2’s extrastriate areas are also largely intact (Figure S3). Nevertheless, this subject has a dense quadrantanopia (Figure 2). One possible explanation may be that the level of activity elicited by visual stimulation in this patient is not sufficient to support useful vision. However, the mean amplitude of pRFs covering the scotoma in subject P2 is decreased by only ~27% compared to “seeing” locations, both in area V1 and hV5/MT+. It is unlikely that this could be the sole cause for the perceptual defect, since: 1) such decreases are routine when presenting stimuli of low contrast without affecting visual perception, 2) similarly low pRF amplitudes sometimes occur in seeing locations (area hV5/MT+ of patient P4). These observations suggest that the BOLD signal amplitude of the pRF maps, a surrogate measure of visual modulation strength, is not necessarily a good indicator of residual visual perceptual capacity in subjects with cortical lesions. Instead, disrupted or poorly synchronized organization of visual processing, which is not directly measured by the BOLD signal, is likely to play a significant role in the loss of visual perception.

### Responses arising from V1-bypassing pathways

For 2/5 patients (P1, P4) there are visual field regions covered by area hV5/MT+ that are not covered by V1. In fact, the part of V1 corresponding to these areas of the visual field is anatomically lesioned. Since activity in these locations arises from stimuli presented well within the scotoma (Figure 3), this strongly suggests that there are V1-bypassing pathways capable of activating area hV5/MT+ in these patients. Approximately 50% (13/27) of visually responsive voxels in hV5/MT+ of patient P4 could not be explained by activity in area V1 during cortico-cortical receptive field mapping, suggesting the existence of V1-bypassing pathways. By contrast in AS controls, essentially all hV5/MT+ voxels had associated well-defined, retinotopically corresponding, CFs. All hV5/MT+ voxels with pRFs inside the scotoma for patient P1 and a few (~5) hV5/MT+ voxels with pRFs inside the scotoma for patient P4, were linked by CF modeling to V1 voxels with pRFs lying in the inferior (seeing) quadrant. This suggests that these hV5/MT+ voxels do receive part of their input ectopically from spared V1 voxels. However, hV5/MT+ voxels with pRFs inside the scotoma also show responses when the visual stimulus is well inside the blind quadrant (Figure 3), which are unlikely to arise from these ectopic V1 locations with pRFs inside the seeing quadrant, since V1 pRFs are small and do not extend to the required visual field locations. This again argues that responses in these hV5/MT+ voxels within the scotoma arise at least in part from V1-bypassing pathways. Activity in these pathways however is not sufficient to reduce the size of the scotoma measured in visual field perimetry.

### Visual awareness associated with activity in hV5/MT+

In contrast to prior reports (Barbur et al., 1993; Zeki and Bartels, 1999) our results suggest that activation of hV5/MT+ alone is not sufficient for visual awareness (Goebel et al., 2001; Barleben et al., 2015). Other studies have pointed out the importance of feedback projections from V5/MT to V1 (Cowey and Walsh, 2000; Pascual-Leone and Walsh, 2001; Silvanto et al., 2005) supporting the idea that V1 is critical for conscious perception. Here we showed that V1 and hV5/MT+ can both show visually driven BOLD activity, but this is not necessarily a sufficient condition for conscious perception. It is unlikely that the lesion has selectively damaged the feedback connections from hV5/MT+ to V1 while sparing feedforward connections, and in particular for subject P2 that had an optic radiation lesion, early visual areas remain intact. Our results suggest two possibilities that can contribute to the loss of visual perception in the presence of residual area V1 BOLD activity: 1) Activity across visual areas becomes too asynchronous or disorganized for visual stimulus awareness to arise (Pollen, 1999; Silvanto, 2015), but this is not directly reflected in the BOLD signal. 2) In some cases, extrastriate visual areas V2/V3 play an important role in visual awareness (Horton and Hoyt, 1991; Merigan et al., 1993; Slotnick and Moo, 2003; Salminen-Vaparanta et al., 2012) and their injury contributes significantly to the loss of visual perception. However, (2) is not the case for patient P2 whose early visual areas are intact and show activity corresponding to the visual field scotoma.

An important point to reinforce here, in agreement to Papanikolaou et al. (2015), is that fMRI pRF measurements provide information about the underlying pathophysiology of the visual field scotoma not necessarily revealed by standard methods of visual field perimetry. The potential caveat should be mentioned here, that the stimuli used to assess the perceptual visual field scotoma (static and kinetic perimetry tests) differ from the visual stimulus that was used to map pRF boundaries. However, i) patients did not report seeing the visual stimulus we used to map their areas by fMRI in their scotomatous field, and ii) the depth (<-20dB) of the visual field scotoma measured in Humphrey’s perimetry is too large to readily explain the extent and strength of visual modulation seen within the scotoma by fMRI. It remains an open question, well worth pursuing, whether the development of new perimetry methods may better reflect the different underlying visual area activation pathophysiology phenotypes revealed by fMRI.

Visual-motion related activity in area hV5/MT+ has been associated with subconscious visual perception in patients with V1 lesions, a phenomenon called “blindsight”. Studies in humans and animal models suggest that blindsight is mediated by subcortical pathways, which effectively bypass area V1 and transmit information from the retina to extrastriate visual cortex (Weiskrantz et al., 1977; Weiskrantz, 1996; Stoerig and Cowey, 1997). Another possible explanation, for some cases, is that there are spared functional V1 “islands” that may mediate residual vision (Gazzaniga et al., 1994; Morland et al., 2004; Radoeva et al., 2008). Our patients have not been tested for blindsight performance, but their performance within the scotoma in a motion direction discrimination task was at chance for 3 patients examined (P1, P2, P5). It would be interesting in future research to investigate whether the different patterns of activation observed in hV5/MT+ are associated with blindsight performance.

### Reorganization of hV5/MT+ following V1 injury

We found increased pRF sizes in hV5/MT+ of the lesioned hemisphere for three out of five patients (P1, P3, P5) compared with AS controls, suggesting reorganization. The increase in pRF size may occur because subcortical inputs from LGN or the pulvinar reorganize via sprouting of cortical axons and contribute to hV5/MT+ activation. Interestingly, the same patients had increased pRF size in hV5/MT+ of the contra-lesional (healthy) hemisphere. It is therefore possible that the pRF size increase occurs as a result of reorganization of callosal connections between the two hV5/MT+ complexes (Silvanto et al., 2007; Bridge et al., 2008). Patients P2 and P4 had a similar pRF size in hV5/MT+ of the lesioned hemisphere as the AS controls. These patients had decreased representation in hV5/MT+ of the contra-lesional hemisphere and particularly patient P2 had significantly smaller pRF sizes, suggesting that the pathways to hV5/MT+ of the healthy hemisphere in these patients are affected by the lesion. The differences between patients may arise by the fact that each patient has a unique lesion. Patient P2 for example has an optic radiation lesion and patient P4 has a larger V1 lesion compared to the other patients that may have affected the capacity for reorganization in hV5/MT+ of these patients. More studies with a larger number of patients will be needed in the future to improve our understanding of visual processing in the context of injury.

### Conclusions

We found that pRF maps of ipsi-lesional area hV5/MT+ overlap significantly with regions of the dense perimetric scotoma. In some subjects, hV5/MT+ responses appear to be mediated by spared V1 to hV5/MT+ projections while, in others, by V1-bypassing pathways or a combination of both. Apparently, hV5/MT+ activation, even if it is accompanied by activity in corresponding parts of V1 cannot guarantee visual perception. Interestingly, fMRI measurements provide information about the underlying pathophysiology of the visual field loss not necessarily revealed by standard methods of visual field perimetry. A timely question to ask is whether regions of the scotoma have different capacity for recovery based on their profile of coverage by spared visual areas. If this turns out to be the case, a better characterization of visual field coverage maps in spared visual areas following visual cortex lesions could help us develop better strategies for visual rehabilitation (Papageorgiou et al., 2014; Smirnakis, 2016).

## Acknowledgements

This work was supported by the Max Planck Society, the Deutsche Forschungsgemeinschaft (DFG), the Plasticise Consortium (Project HEALTH-F2–2009–223524), National Eye Institute R01 (EY024019) to SMS, and a Postdoctoral Ruth L. Kirschstein National Research Service Award, NIH to TDP. AP is supported by a DFG Research Fellowship. We thank Elke Krapp, Eleni Papageorgiou, Katarina Stingl and Anna Bruckmann for their help with patient recruitment and MRI scanning and Natalia Zaretskaya for her help with the eye-tracking. We thank Koen Haak for providing us with the MATLAB toolbox for running the connective field modeling method. The authors declare no competing financial interests.

